# Mammalian Milk Glycomes: Connecting the Dots between Evolutionary Conservation and Biosynthetic Pathways

**DOI:** 10.1101/2023.02.04.527106

**Authors:** Luc Thomès, Viktoria Karlsson, Jon Lundstrøm, Daniel Bojar

## Abstract

Milk oligosaccharides (MOs) are among the most abundant constituents of breast milk and are essential for health and development. Biosynthesized from monosaccharides into complex sequences, MOs differ considerably between taxonomic groups. Even human MO biosynthesis is insufficiently understood, hampering evolutionary and functional analyses. Using a comprehensive resource of all published MOs from >100 mammals, we develop a nonparametric pipeline for generating and analyzing MO biosynthetic networks, which readily generalizes to other glycan classes. We then use evolutionary relationships and inferred intermediates of these networks to discover (i) distributional glycome biases, (ii) biosynthetic restrictions, such as reaction path dependence, and (iii) conserved biosynthetic modules. This allows us to prune and pinpoint biosynthetic pathways despite missing information. Machine learning and network analysis cluster species by their milk glycome, identifying characteristic sequence relationships and evolutionary gains/losses of motifs, MOs, and biosynthetic modules. These resources and analyses will advance our understanding of glycan biosynthesis and the evolution of breast milk.

## Introduction

Breast milk is a complex mixture of highly functionalized molecules, produced by all mammals^1^. Complex carbohydrates, or glycans, are among the most abundant components of breast milk^2^. In particular, free milk oligosaccharides (MOs), secreted by the mammary gland, are abundant and exhibit a range of nutritional and defensive properties^3,4^. Comprising chains of monosaccharides arranged in defined linkages and forming branches, the structural diversity of MOs is high. Anti-pathogen functions of MOs usually hinge on terminal motifs (sialylation, fucosylation, etc.)^5,6^ that can serve as binding epitopes, indicating the importance of the motif repertoire for MO function^3^.

The canonical biosynthesis of MOs starts from lactose and, in a strictly stepwise manner, adds monosaccharide via glycosyltransferase enzymes to elongate or branch the eventual MO^4,7^. Glycosyltransferases are sensitive to (i) the identity of the transferred monosaccharide, (ii) the sequence context to which they add the monosaccharide, and (iii) the orientation in which the monosaccharide is added^8^. The difficulty of assigning enzymes for the biosynthesis of a specific structure lies in the existence of closely related enzyme (e.g., eight iso-enzymes of the B3GNT family in mammals, with differing specificities), and in their action on one or several glycan classes (e.g., glycolipids, protein-linked glycans, glycosaminoglycans, etc.).

While MOs are universal in mammals, substantial inter-species differences in MO repertoires have been noted^9^, e.g., a presumed paucity in aquatic mammals^10^, a particular complexity in humans^11^, or the usage of exotically modified monosaccharides in monotremes^12^. However, while comparisons within^13,14^ and between^10,15–18^ species exist, a comprehensive overview across Mammalia is still lacking. Further, existing comparisons usually contrast individual structures or manually selected motifs, lacking the rigor of state-of-the-art computational analyses and neglecting the biosynthetic dependencies of MOs. This has led to an incomplete understanding of the evolution of MOs^9^ and, consequently, of breast milk.

Computational analyses of the biosynthesis of the (milk) glycome usually start with a repertoire of general enzyme activities explaining observed MOs and grow the biosynthetic network based on this^7,19,20^. This has been used to predict yet-to-be-discovered MOs and understand MO biosynthesis. However, this approach often results in networks that are simultaneously too permissive, with up to millions of possible structures, and too restrictive, disallowing reactions not specified beforehand. To facilitate this, most approaches only consider a subset of the milk glycome that is explainable by known enzyme activities and tend to exclude rare or noncanonical structures. Further, previous approaches focused predominantly on human MOs, forestalling evolutionary inferences and the usage of conservation as a proxy for function. In general, the necessity of specifying operating enzyme activities and/or even rate constants strongly limit the generalizability of previous approaches for different glycan classes or taxonomic species, as well as their usefulness for further elucidating biosynthesis beyond known facts.

Here, we use a comprehensive, curated dataset of ~2,300 literature MO-species associations from >100 species, including recent extensive experimental data from nine new species^18^, leveraging evolutionary relationships and the connectedness of MO biosynthesis to construct and analyze biosynthetic networks. We present a generalizable approach to establish evolution-informed biosynthetic networks of glycan types, such as MOs, without requisite knowledge about enzymes or enzymatic functions. Integrated into the popular glycowork package, this allows researchers to construct glycan biosynthetic networks with a single line of code by providing a list of glycans, regardless of which glycan class they stem from. For the example of MOs, we then engage in analyzing these biosynthetic networks and find systematic biases in unobserved intermediaries, conserved reaction path dependencies across species and sequence contexts, and conserved biosynthetic modules. We further reveal likely losses and gains of these modules in the evolution of Mammalia. Our findings shed light on glycan biosynthesis coordination in general, the conserved constraints of MO biosynthesis in particular, and the evolution of breast milk across Mammalia.

## Results

### Constructing biosynthetic networks of the free milk glycome

Solely analyzing the properties of individual MOs would belie their shared biosynthetic history. Given that MOs originate in their biosynthesis from, at maximum, two “roots” (lactose and lactosamine), they can be conceptualized as a network with, at maximum, two connected components. Each node of this network would be a MO and each connecting edge a biosynthetic step, adding exactly one monosaccharide or post-biosynthetic modification, such as sulfation. However, not all intermediate MOs may be observed with mass spectrometry or related techniques, due to low abundance for instance. Additionally, the identity of the involved enzymes is still incompletely understood, even in humans^4,7^. Therefore, we developed an empirical algorithm that does not rely on known enzymatic activities and that finds the minimum number of proposed intermediates to connect all observed structures in a biosynthetic network (see STAR Methods), conceptually vaguely related to the Compozitor framework^21^, yet not limited to compositions but also taking glycan structure into account. In this, glycans are conceptualized as nodes, connected by edges representing biosynthetic steps.

We developed this process into a general analysis platform, added as the *network* module to the open-access Python package glycowork^22^ for the analysis of glycans and their role in biology (Figure 1A), to facilitate other inquiries into glycan biosynthetic networks. Given a list of glycans, functions within this module can flexibly construct, prune, and analyze biosynthetic networks at scale on a personal computer, usually in seconds to minutes. The *construct network* function automatically detects glycan class and thus generalizes readily to other glycan classes, such as *O*-linked glycans on proteins or glycolipids (Figure S1).

**Figure 1.**
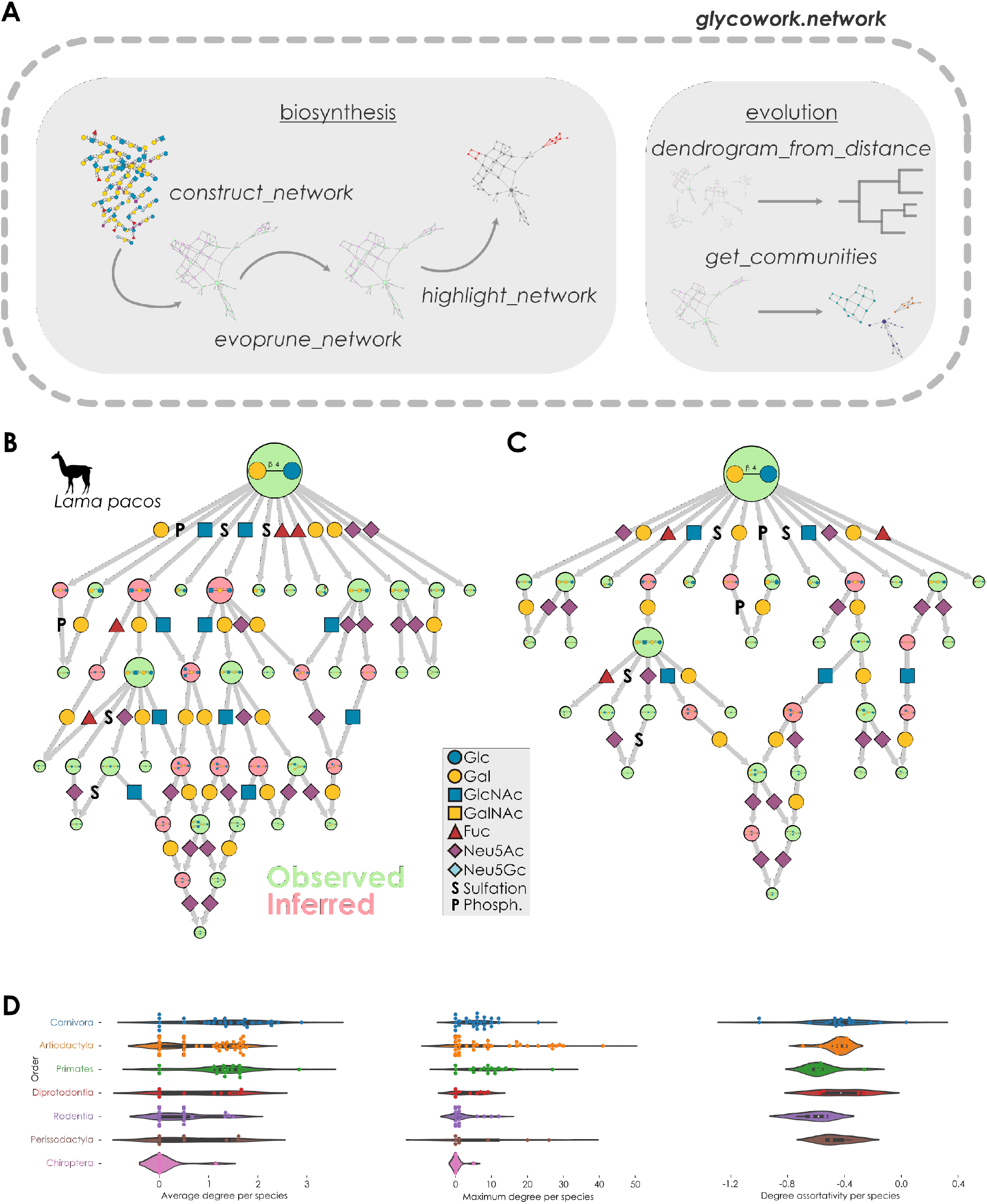
Species-specific biosynthetic networks of MOs. **(A)** Schematic view of the network module of glycowork. This module contains all relevant code to construct and analyze glycan biosynthetic networks. Example workflows are shown for the two submodules, biosynthesis and evolution. (**B**) Biosynthetic network of alpaca milk glycans. Each node depicts a milk glycan (green: observed, pink: inferred), connected by directed edges symbolizing biosynthetic steps (coded by the Symbol Nomenclature for Glycans, SNFG). Node size is scaled by out-degree. (**C**) Biosynthetic network of alpaca milk glycans after pruning. We used the network from (B) and removed inferred nodes from paths in diamond-shaped network motifs using the *evoprune_network* function in glycowork with default parameters (see STAR Methods for details). All networks were visualized with Cytoscape (version 3.9.1), although they can also be easily visualized using the *plot_network* function in glycowork. (**D**) For pruned networks from all species, we calculated their average degree, maximum degree, and degree assortativity. Only taxonomic orders with at least five species were included.

We first constructed a biosynthetic network from the recently measured alpaca MOs (*Lama pacos*, Figure 1B). This network clearly showed preferences during MO biosynthesis, with inner reactions that were dominated by the addition of galactose (Gal) and *N*-acetylglucosamine (GlcNAc) and terminal reactions with an enrichment for fucosylation and sialylation, monosaccharides usually adorning non-reducing ends of mature MOs^23^. Proposed intermediates offered paths to particularly elongated MOs.

While inferred intermediates yield a connected network, they also introduce redundancies, as several paths might connect to an observed MO^19^. We developed “evolutionary pruning” to ameliorate this redundancy by leveraging evolutionary information. Briefly, we extracted diamond-shaped subnetworks of two glycans connected via two possible intermediates (reactions A-then-B versus B-then-A, resulting in two different intermediate structures). Then, we followed by assessing how often each “path” is taken in Mammalia, using the >100 species in our dataset. Overall, both network construction and pruning scaled well with typically encountered sizes of glycan classes (Figure S2). In pruning, based on a probability threshold (usually 0.01 here, for maximum stringency), we removed unlikely paths from the network (see STAR Methods). On average, our biosynthetic networks contained 26.1% inferred nodes, of which we pruned approximately a quarter at the chosen threshold. Pruning the alpaca network resulted in a more modular network and higher confidence in remaining predicted intermediates (Figure 1C). Subsequent analyses relied on these pruned networks.

Next, we wanted to gain an overview of network properties across our dataset. We compared the average degree, maximum degree, and degree assortativity of biosynthetic networks of all our species (Figure 1D). The degree of a node is its number of connections, which can be averaged over the whole network. Degree assortativity is the correlation of degrees of neighboring nodes (i.e., whether nodes connect to similarly connected nodes). Species in the well-represented orders Primates, Artiodactyla, and Carnivora exhibited similar average node degrees, yet differed in their maximum node degrees. Artiodactyla species seemed to have a higher propensity for highly connected hub nodes. Species in all orders exhibited negative degree assortativity, implying that hub nodes were connected to sparsely connected nodes.

### Biosynthetic constraints in the free milk glycome

We next investigated why some inferred nodes were not experimentally measured. This also revealed that some MOs were almost exclusively experimentally observed (Figure 2A). These MOs, structures such as 2’-fucosyllactose (2’-FL) or 3’sialyllactose (3’-SL), constituted biosynthetic “dead-ends” that were barely elongated, if at all. Conversely, we noted that a second cluster of glycans almost always were inferred intermediates. A common element of these unobserved MOs was that their non-reducing ends tended to terminate in GlcNAc, often as an “unfinished” LacNAc repeat (Galβ1-4GlcNAc), also noted previously^19^. The absence of these intermediates – which must exist to produce the observed structures – from measurements could indicate coordinated action by GlcNAc transferases and galactosyltransferases, yielding low concentrations of GlcNAc-terminated intermediates. In general, inferred intermediates exhibited higher connectivity than measured glycans (Figure 2B), potentially indicating their rapid metabolization by a greater number of available pathways. These findings were supported by enrichment analyses of terminal motifs (Figure 2C), showing that inferred intermediates were highly enriched in all GlcNAc linkages and Fucα1-2/3/4, suggesting that fucosylation aggravates the absence of GlcNAc-terminated glycans. Observed structures, on the other hand, were primarily enriched in terminal sialic acid, galactose, and GalNAc (Figure 2D), supporting our findings of biosynthetic “dead-ends” and finished LacNAc repeats. These observations stress the relevance of applying a systems-level perspective to this topic.

**Figure 2.**
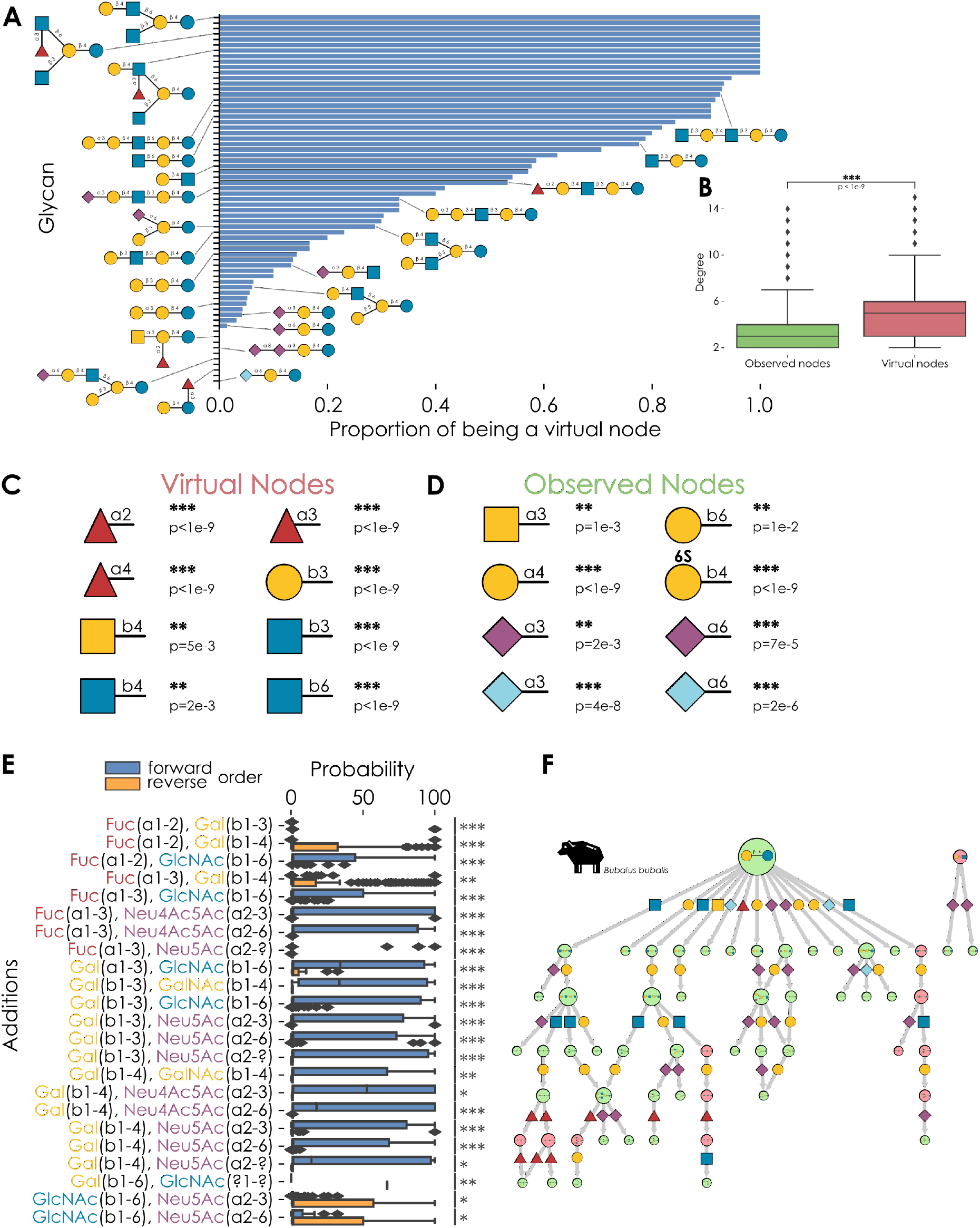
Biosynthetic constraints in MOs. **(A)** Propensity of common MOs to be inferred or observed. For glycans in networks of at least 10 species, we assessed their proportion as an experimentally determined glycan or an inferred intermediary. Glycan structures are shown via the SNFG. (**B**) Propensity of virtual nodes being network hubs. Excluding leaf nodes, we used a one-tailed Welch’s t-test to assess whether predicted intermediates exhibited a higher degree in pruned biosynthetic networks than observed structures. (**C-D**) Enrichment analysis of terminal motifs in virtual (C) and observed (D) nodes. For all terminal motifs, statistical enrichment was tested via one-tailed Welch’s t-tests with a Holm-Šídák correction for multiple testing. (**E**) Biosynthetic reactions with path dependence. For each observed addition of two monosaccharides in diamond-shaped network motifs, we grouped their probabilities across glycan sequences (i.e., monosaccharides A and B could be added to glycan X, Y, Z, etc.) in both the forward (A-then-B) and reverse (B-then-A) order. Shown are additions with a mean difference of at least 15 occurring in more than two sequences. Mean difference was tested via two-tailed Welch’s t-tests. Data are depicted as mean values, with box edges indicating quartiles and whiskers indicating the remaining data distribution. ***, p < 0.001; **, p < 0.01; *, p < 0.05; n.s., p > 0.05. (**F**) Maximum-likelihood biosynthetic network for *Bubalus bubalis* MOs. Based on the reaction path probabilities from Figure 2E, we assessed all remaining diamond-shape motifs in the pruned network and chose a maximum-likelihood path if (i) a two-sided Welch’s t-test exhibited a significant difference in path likelihood (p < 0.01) and (ii) the removal of the alternative path did not abolish network connectivity.

The fact that some inferred nodes comprised a path that was never taken and could be safely pruned with our evopruning approach (Figure 1C), implied the path dependence of some biosynthetic reactions. To investigate this systematically, we analyzed all instances where two biosynthetic paths led to the same MO. As the individual enzyme activities for each step in both paths must exist to create the final MO, network inferences in previous work routinely proposed both paths. However, enzymes may not add monosaccharides in any order. We thus hypothesized that one path would be at least biosynthetically favored, if not exclusively used.

Using the evolutionary information in our multi-species biosynthetic networks, we calculated, for each pair of non-contiguous monosaccharide additions, the probability of A-then-B and B-then-A (see STAR Methods). This mostly resulted in substantially unequal distributions, with clear preference of one order of addition (Figure 2E), indicating that reaction path dependence is conserved across species and glycan contexts (as the addition of two monosaccharides can be monitored on different “backbones”). We stress that we here only refer to non-contiguous reactions, namely reactions in which the two added monosaccharides are not added in direct succession/connection. Path dependence of non-contiguous addition also seems conserved across reaction families, e.g., with galactose being added before Neu5Ac across different linkages of either monosaccharide. Simplifying, we observed a reaction order of Gal > Fuc ~ GalNAc > Neu5Ac > GlcNAc. Some reaction pairs, such as Neu5Acα2-3 and Neu5Acα2-6, did not exhibit significant path dependence (Figure S3). We also identified cases in which the reaction path dependence showed a divergent pattern across different mammalian orders (Figure S4). Species in Artiodactyla were for example unique in adding Galβ1-3 and Neu5Acα2-3 via both Gal-then-Neu5Ac and Neu5Ac-then-Gal, highlighting the existence of at least some unique reaction path dependences across taxonomic orders. We note that Neu5Ac-containing paths in general were particularly prone to exhibiting order-specific reaction dependence, which reached statistical significance (p = 0.011, one-sided Welch’s t-test of path preference standard deviation of Neu5Ac-containing paths across taxonomic orders versus other paths).

We then further specified biosynthetic networks by choosing the maximum-likelihood paths based on biosynthetic constraints and reaction path dependence (Figure S5), by choosing the path with a significantly higher reaction path probability (see STAR Methods). For networks such as from buffalo milk, this yielded a nearly fully-specified network (Figure 2F). Repeating this for all species yielded high-confidence inferred intermediates (Table S2) and a robust basis for later investigations.

### Identifying conserved biosynthetic modules for the free milk glycome

The reaction order preference for most monosaccharides led us to speculate that whole parts of the MO biosynthetic network are conserved, also seen by the increasing modularization of the network upon evolutionary pruning (Figure 1C) and reaction path choice (Figure 2F). This was reinforced by for instance observing a visual separation of fucosylated and sialylated glycans in biosynthetic networks, which we further investigated and revealed that most MOs are either fucosylated or sialylated, rarely both (Figure 3A), with seeming biosynthetic trajectories that are locked-in early. This indicated carefully orchestrated biosynthesis modules, resulting in defined biochemical populations with distinct motifs.

**Figure 3.**
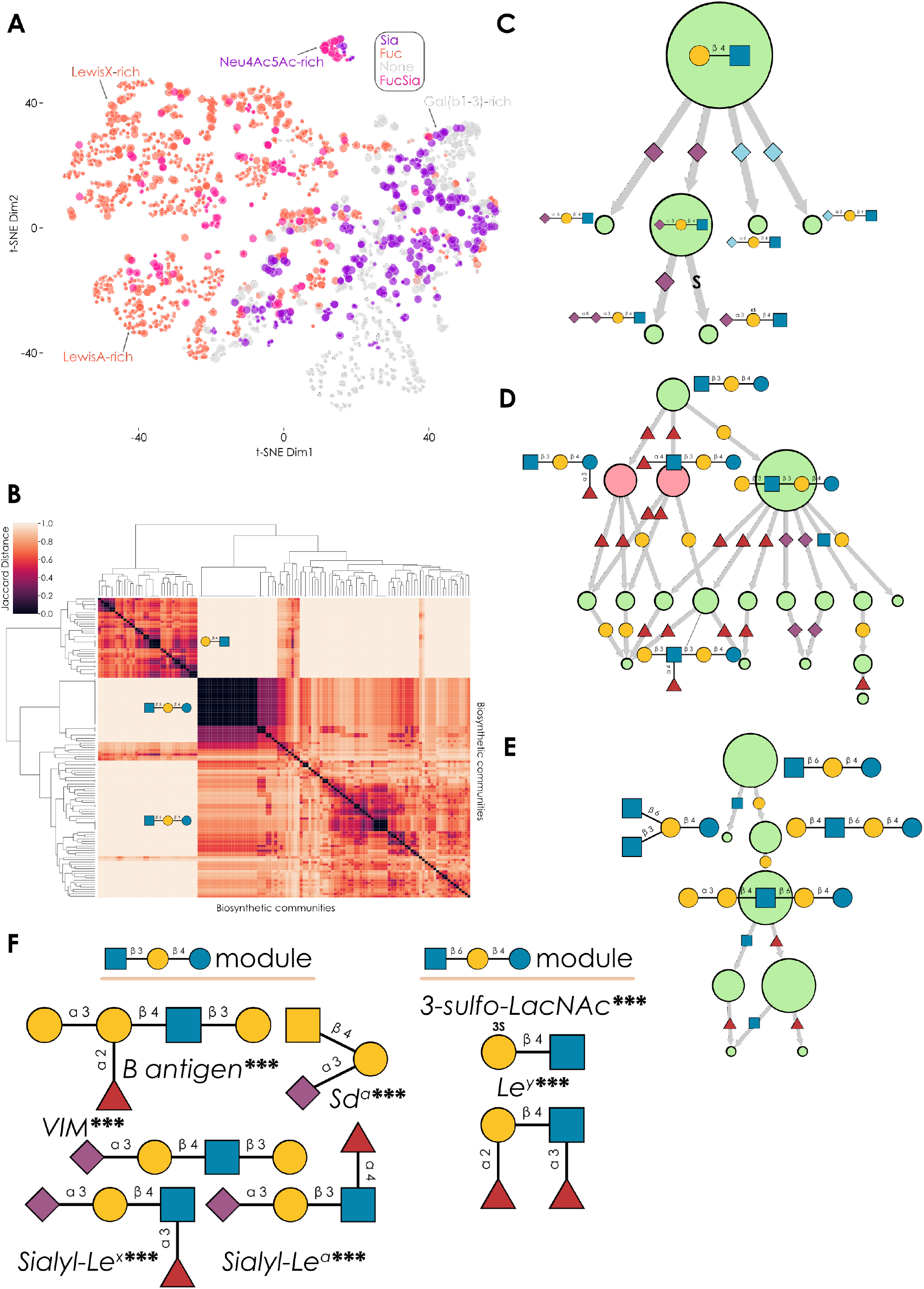
Conserved biosynthetic modules in milk oligosaccharide synthesis. **(A)** Learned representation of milk glycans. Using a SweetNet-type model trained on predicting taxonomic species from glycans^5^, we retrieved representations of all unique milk glycans in our dataset via the *glycans_to_emb* function from glycowork, depicted them via t-SNE, and colored them as to their fucosylation and sialylation status. Node size is scaled by probability of being experimentally observed versus inferred, across our set of species. (**B**) Hierarchical clustering of biosynthetic modules. For all networks, we extracted biosynthetic modules via the Louvain community detection algorithm, calculated their pairwise Jaccard distances, and clustered them, excluding communities only consisting of lactose. (**C-E**) Conserved biosynthetic modules. We used density-based spatial clustering of applications with noise (DBSCAN) to extract conserved biosynthetic modules from communities in (B), with examples shown. (**F**) Enrichment analysis of terminal motifs in glycans from the GlcNAcβ1-3 and GlcNAcβ1-6 modules. For all terminal motifs, statistical enrichment was tested via one-tailed Welch’s t-tests with a Holm-Šídák correction for multiple testing. Representative enrichments that are not observed in the other module are shown. ***, p < 0.001; **, p < 0.01; *, p < 0.05; n.s., p > 0.05.

To systematically identify these sub-networks, we used the Louvain method^24^ to extract communities, sub-networks that are densely connected internally but sparsely connected to the rest of the network. This yielded several communities that formed clusters with low Jaccard distances (i.e., high network similarity; Figure 3B). Using density-based clustering, we retrieved the most conserved biosynthetic modules, comprising three modules: (i) lactosamine-based MOs (Figure 3C), (ii) MOs derived from the progenitor GlcNAcβ1-3Galβ1-4Glc (Figure 3D), and (iii) MOs starting from the progenitor GlcNAcβ1-6Galβ1-4Glc (Figure 3E). For orientation, we note that further extension of (ii) and (iii) corresponds to LNT/LNnT and iso-LNT/iso-LNnT, respectively.

These modules had substantial consequences for the motifs eventually displayed on “mature” MOs. Statistical enrichment/depletion in the GlcNAcβ1-3 and the GlcNAcβ1-6 modules (Figure 5F), uncovered motifs such as Sd^a^ or sialyl-Lewis structures to be only enriched in the GlcNAcβ1-3 module, while the GlcNAcβ1-6 module was enriched for sulfated structures and the Le^y^ motif. We also noted an approximately equal distribution of the modules in lactosamine-based MOs (37% GlcNAcβ1-3, versus 39% for GlcNAcβ1-6). While some interconversion does exist, most notably in structures, such as LNnH, that exhibit both GlcNAcβ1-3 and GlcNAcβ1-6 branches, the two modules are largely separated. The early position of these module progenitors in the network implies that biosynthesis is “locked-in” early, also shown by the distinctiveness of the resulting modules, enabling the network to explore different regions of biosynthetic space.

### Tracing the evolution of the free milk glycome

As both individual reactions and biosynthetic modules seemed conserved across wide swaths of Mammalia, we hypothesized that the resulting milk glycome, its MOs, and their motifs, would also be conserved. Clustering mammalian species according to the similarity of their milk glycomes (Figure 4A) yielded clusters that broadly corresponded to taxonomic orders, especially considering the vastly different degrees of coverage, with the knowledge about species ranging from single-digit to hundreds of characterized MOs. Prominent among these are the two marsupial orders that cluster together, displaying characteristically different MOs further explored below. We also note that the nine species characterized in our previous work^18^ are found in the same broad clade, despite comprising multiple orders. The multitude of new structures and motifs identified in our in-depth approach makes them distinct from most previous MO descriptions. We therefore believe that a careful re-investigation of already measured milk glycans is likely to uncover greater milk glycome complexity than previously assumed.

**Figure 4.**
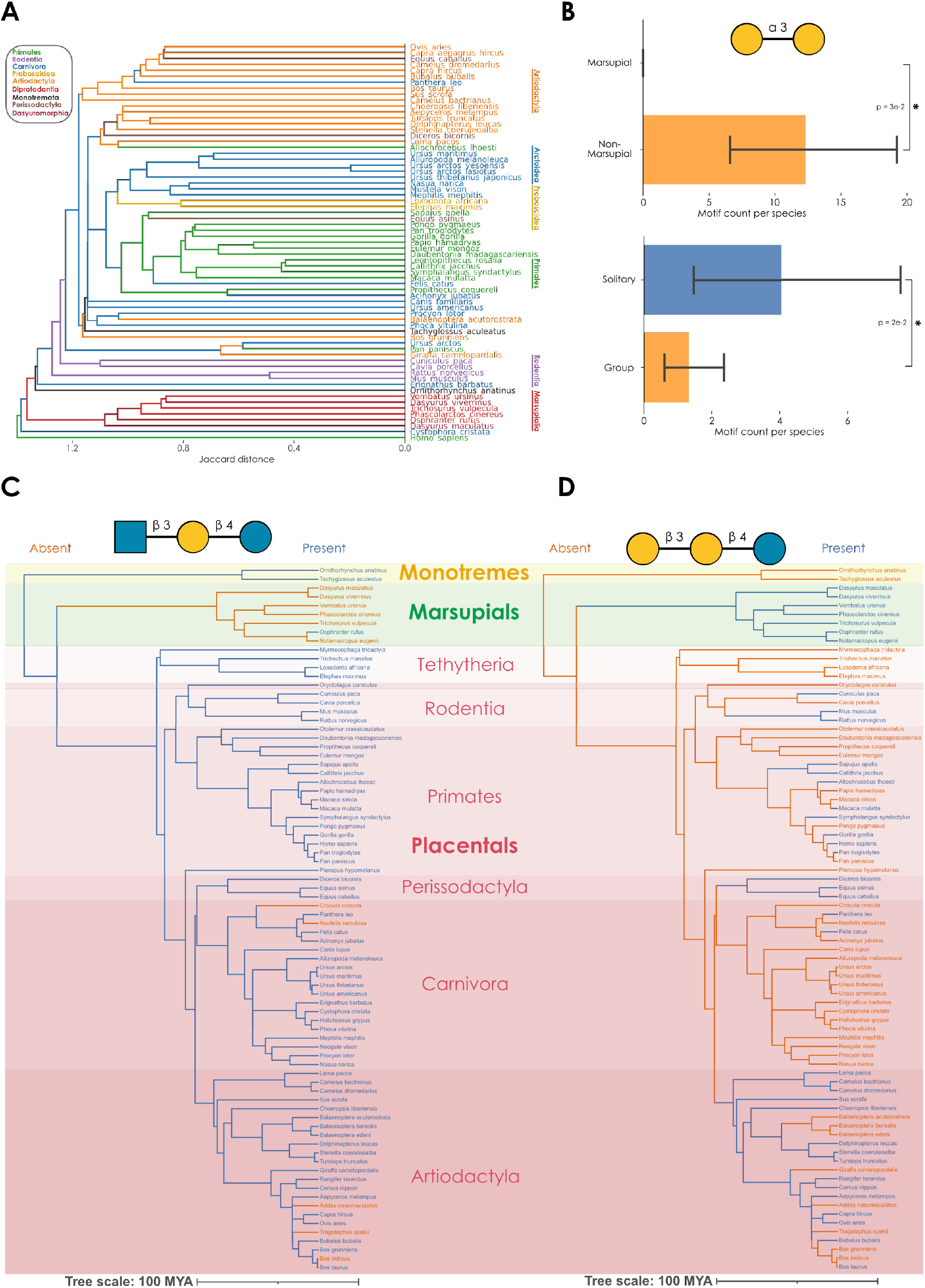
Evolution of the free milk glycome. **(A)** Milk glycome similarity across species. For all species with at least five MOs, we used their pruned biosynthetic network to calculate pairwise Jaccard distances and create a dendrogram via hierarchical clustering. (**B**) Evolutionary conservation of the alpha-Gal motif in MOs. For all species with at least fifty MOs in their biosynthetic networks (inferred or observed), we counted the occurrence of the alpha-Gal motif in their MOs and contrasted this abundance between marsupial / non-marsupial and solitary / group species. Significance was established by one-sided Welch’s t-tests. Data are depicted as mean with 95% confidence interval. ***, p < 0.001; **, p < 0.01; *, p < 0.05 (**C**) Evolutionary loss of GlcNAcβ1-3Galβ1-4Glc milk glycans. For all species with at least five glycans in their pruned networks, we assessed the presence of GlcNAcβ1-3Galβ1-4Glc. Then, we overlayed this information on the evolutionary history of these species, derived from TimeTree^25^. (**D**) Evolutionary gain of Galβ1-3Galβ1-4Glc milk glycans. We used the same procedure as in Fig. 4C to show the gain of this sequence in marsupials.

Given our comprehensive MO dataset, we developed a workflow assessing the evolutionary conservation of a glycan or motif. Using pruned networks, we for instance revealed that lactosamine-based MOs were strongly conserved in even-toed and odd-toed ungulates (Artiodactyla and Perissodactyla) and considerably rarer in many other mammalian orders (Figure S6). Further, motifs such as the alpha-Gal motif (Galα1-3Gal), absent in several groups such as Old World primates^26^ or marsupials (Figure 4A), were generally depleted in non-solitary mammals (Figure 4B). This showcases glycomic environmental adaptation across evolutionary similarity^5,27^. We further hypothesized that glycans that are more central to the network would also be more likely to be conserved in a given taxonomic order and indeed found a moderate correlation between degree of conservation and betweenness centrality (i.e., importance for network connectivity) in the order Diprotodontia (Spearman rank correlation; rs = 0.4, p = 0.003; Fig. S7).

The separate grouping of marsupials (Figure 4A) led us to hypothesize that their conserved biosynthetic modules also differed. We thus analyzed the presence of a module progenitor, GlcNAcβ1-3Galβ1-4Glc, in the biosynthetic networks of all species. The benefit of this approach is that this also considers species that do not have a published GlcNAcβ1-3Galβ1-4Glc sequence but still have this glycan as an inferred node that is central to network connectivity, making these inherently incomplete data more comparable. Combining this information with species divergence data uncovered that GlcNAcβ1-3Galβ1-4Glc was absent from all marsupials except for *Osphranter rufus* and present in nearly all placentals (Figure 4C). It should be noted that all mentions of the GlcNAcβ1-3Galβ1-4Glc module in *O. rufus* originate from a single publication^10^ using a C18 column in reverse-phase chromatography that cannot confidently distinguish between GlcNAcβ1-3 and GlcNAcβ1-6 isomers, likely indicating a misannotation and the complete loss of this module in marsupials. The absence of the GlcNAcβ1-3Gal unit from marsupial MOs has also been pointed out before^9^. As we also find this progenitor MO, and its consequent biosynthetic module, in monotremes, we conclude that marsupials have lost this biosynthetic module at some point after the divergence of Monotremata from the rest of Theria, approximately 166 million years ago^28^. Interestingly, marsupial genomes still contain the enzyme purported to catalyze this reaction, B3GNT2^7^, and only lack B3GNT6 - the enzyme responsible for synthesizing the core 3 structure in *O*-glycans - from the B3GNT family.

Conversely, we found the evolutionary gain of a module that is highly enriched in marsupials (Figure 4D) and potentially in some parts of Artiodactyla. The progenitor of this module, Galβ1-3Galβ1-4Glc, uses the same starting point as the GlcNAcβ1-3 module (C3 of lactose), directly competing with this module for substrates, and potentially constituting a replacement for the loss of the GlcNAcβ1-3 module in marsupials. This also could have resulted in the characteristic marsupial oligo-galactose MOs^9^, which start from this progenitor. Additionally, this new module in marsupials is the cause for the relatively unique presentation of 3S-modified MOs^18^. Notably, the combination of our rich evolutionary dataset and flexible network methodology can be used to easily engage in similar analyses for other modules or motifs. We thus conclude that our presented data and method platform can be used to investigate various facets of the evolution of breast milk and its constituents.

## Discussion

Despite fulfilling roles from nutrition^29^ to immunity^30^, the biosynthetic space of MOs is not evenly accessible to all mammals, as shown here. With various systems-level approaches, and leveraging evolutionary information, we focused on overcoming the challenge of incomplete information in analyzing the milk glycome, e.g., due to unobserved intermediates. Our networks also became increasingly modular with our pruning strategies, emphasizing the benefits of this approach versus analyzing all possible alternatives. This strategy unveiled broad evolutionary conservation of motifs, glycans, reaction path dependence, and biosynthetic modules. Our broad dataset and approach enabled us to pinpoint evolutionary losses and gains in mammalian MO biosynthesis. With this established, we envision further functional inquiries into these evolutionary changes.

A limitation could be that we average out several effects in breast milk. Colostrum, on average, has higher MO diversity than mature milk^31^. Further, even mature milk differs in composition during lactation^31^. Yet the academic literature presents a mixture of everything currently known, often without information of milk age. Our dataset represents overall species capabilities, while disentangling temporal effects on biosynthetic networks will require further research. Future avenues could also consider glycan abundance – as two species might have identical MO repertoires but direct their metabolic flux into different directions – or intra-species diversity^13,14^. We do note, however, that the absence/presence type of information we use here to construct biosynthetic networks already results in largely accurate clustering of taxonomic orders. Finally, despite the efforts of about a century of milk oligosaccharide research, large fractions of the pan-mammalian milk glycome remain unexplored and certainly contain yet more surprises.

We also note that the “locking-in” effect into conserved biosynthetic modules, that we observe early on in MO biosynthesis, partly determines the resulting motif repertoire, which could allow for predictive evaluations of which motifs will likely be discovered in future milk glycomics studies. One example can be found in the recently discovered LacdiNAc motif in MOs^18^. Given that LacdiNAc occurrences seem to be enriched for the GlcNAcβ1-3 biosynthetic module, we expect that marsupials, having lost this module, do rarely exhibit this motif in their MOs, if at all.

It is still an open question why unfinished LacNAc repeats are so rarely experimentally observed. Potential reasons include enzymatic coupling, resulting in lowly abundant intermediate structures, or digestion by microbes. Regardless, we know that these MOs must exist, as no known enzyme transfers Galβ1-4GlcNAc *en bloc* onto glycans. While we tend to favor an explanation of efficient galactosyltransferases and the existence of multiple pathways metabolizing GlcNAc-terminated MOs, due to the network centrality of these structures, that still leaves open the question of whether there is an evolutionary reason to avoid GlcNAc-terminated MOs, e.g., because of instability due to degradation by bacterial exoglycosidases.

An expansion of the data analyzed here would result in even more stringent estimates of reaction path dependencies, rules for unobserved intermediates, and conserved biosynthetic modules. We therefore envision our approach gaining strength as research continues on MOs. We further speculate that similar phenomena and trends might be discovered in other classes of glycans and are confident that the methodology presented here might also provide insight into their constraints and evolution. We thus envision that this approach will advance glycobiology and our understanding of biological processes involving glycans.

## Acknowledgments

This work was funded by a Branco Weiss Fellowship – Society in Science awarded to D.B., by the Knut and Alice Wallenberg Foundation, and the University of Gothenburg, Sweden.

## Author Contributions

D.B. conceived the method. D.B., L.T., and V.K. performed computational analyses. D.B., L.T., and J.L. prepared the figures. All authors wrote and edited the manuscript.

## Declaration of Interests

The authors declare no competing interests.

## STAR Methods

### Resource availability

#### Lead contact

Further information and requests for resources should be directed to and will be fulfilled by the Lead Contact, Daniel Bojar

#### Materials availability

This study did not generate new unique reagents.

#### Data and code availability

All code used and developed here, as well as pre-computed biosynthetic networks, are available at https://github.com/BojarLab/MilkGlycansand/or in the Python package glycowork (at https://github.com/BojarLab/glycowork/tree/dev; already publicly accessible and to be released as version 0.7). Data curated or generated here can be found in the supplementary tables as well as stored as internal datasets within glycowork.

### Method details

#### Dataset construction

For details of the literature curation, see previous work^18^. Briefly, all published milk oligosaccharides from 1898 onwards were gathered from PubMed and associated databases to construct a dataset of 2,296 species-specific MOs from 172 species (with 62 species having only been reported to contain lactose in their breast milk).

All milk oligosaccharides were formatted into IUPAC-condensed nomenclature and paired with their full taxonomical information. All MOs can be found in Table S1 and stored within our Python package, glycowork^22^.

#### Biosynthetic network construction

The main rationale of network construction is a backwards-process to find the minimal number of inferred (“virtual”) intermediates that would connect the observed structures. The root nodes here are always lactose and/or lactosamine, and glycans are treated as molecular graphs. Starting from a list of glycans known to occur in the breast milk of a given species, we first construct an adjacency matrix of glycans that can be directly connected via one biosynthetic step (i.e., the addition/removal of one monosaccharide via a given linkage). We then perform all possible modifications at the non-reducing end(s) of the provided glycans *in silico* (i.e., remove a monosaccharide and linkage), to observe whether any of these “virtual”/unobserved intermediaries would connect two or more observed glycans. These connecting intermediaries are then retained within the network constructed from the adjacency matrix, with each glycan as a node and their biosynthetic steps as edges.

Then, for any glycan not yet connected to the root nodes, an iterative procedure is employed to connect them to the network. At every step, a “shell” of biosynthetic intermediaries is generated from the unconnected glycan (all precursors that are one biosynthetic reaction away) and we attempt to find a connected path to the main network via members of the shell. If unsuccessful, the next shell is generated (precursors that are two reactions away), until the estimated distance to the largest observed precursor is reached. If a path is found, we search for the path(s) with the shortest length and add the corresponding inferred nodes and edges to the network. Then, we search for post-biosynthetic modifications in the glycans of the network (e.g., sulfation, phosphorylation, acetylation) and add the corresponding edges to their un-modified precursors in the network, which are also added as inferred nodes to the network if they are not yet contained therein. Finally, generated leaf nodes that are not experimentally observed are being pruned from the network. The network is converted into a directed graph by only retaining edges that add a monosaccharide or post-biosynthetic modification. All code functionality developed and used here has been integrated as readily usable functions in the *network.biosynthesis* module of the upcoming glycowork version 0.7^22^ (already freely available on the dev branch). Networks are built in NetworkX (version 3.0)^32^ but can be exported into any graph format using the *export network* function of glycowork.

#### Network pruning

Starting from a network, we automatically extract all diamond-shaped motifs (a network motif with four glycans, going from a source to a target via two different paths: path A-then-B and B-then-A) that contain at least one virtual node. Then we use all other milk glycome networks to assess whether either path (i) exists in a network and (ii) is experimentally observed or inferred. By calculating the probability for being experimentally observed, we could remove paths from the network that contained virtual nodes that have never been experimentally observed across our entire dataset. Maximum-likelihood pruning was performed by choosing a path if (i) a two-sided Welch’s t-test of reaction path probabilities exhibited a significant difference in path likelihood (p < 0.01) and (ii) the removal of the alternative path did not abolish network connectivity. Here, reaction path probabilities were determined by the probabilities obtained during evolutionary pruning, shown below. Both pruning procedures resulted in a set of virtual nodes that represent conservative predictions for unobserved yet existing milk oligosaccharides, which can be found in Table S2.

#### Assessing reaction path dependence

From unpruned networks of all species in our dataset, we extract all diamond-shape network motifs. It should be noted that diamond-shape motifs explicitly select for contexts in which the two added monosaccharides are not linked, as there would not be two alternative orders otherwise. Each diamond-shape motif contains two probabilities (A-then-B and B-then-A) for a given glycan context. For each combination *i*, and its associated intermediate, we assess its probability by calculating the ratio:

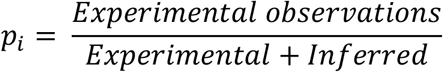

For each unique A-then-B combination, we then average its probabilities across glycan sequence contexts, weighted by how often they occur in different species. We then group matching combinations (i.e., A-then-B with its corresponding B-then-A) to test for significant differences.

#### Extracting and clustering communities in biosynthetic networks

For all species with at least five glycans in their pruned biosynthetic network, we use the Louvain community detection algorithm^24^ from the python-louvain package (version 0.16, https://github.com/taynaud/python-louvain), using the partition of the graph nodes that maximizes modularity. Communities only consisting of lactose are removed. We then calculate pairwise Jaccard distances of communities, defined as the ratio of node set intersection over union. To retrieve clusters, we apply density-based spatial clustering of applications with noise (DBSCAN)^33^ to this distance matrix, using the scikit-learn^34^ (version 1.2.0) implementation with an epsilon of one and a minimum of six samples per cluster.

#### Quantification and statistical analysis

Comparing two groups was done via one-tailed or two-tailed Welch’s t-tests. In all cases, significance was defined as p < 0.05. All multiple testing was corrected with a Holm-Šídák correction. All statistical testing has been done in Python 3.8 using the statsmodels package (version 0.12.2) and the scipy package (version 1.10.0).

## Supplementary Tables

**Supplementary Table 1. Overview of all published milk oligosaccharides.** Related to Figure 1.

**Supplementary Table 2. Milk oligosaccharides including high-probability intermediates.** Related to Figure 2.

## Notes

https://github.com/BojarLab/MilkGlycans

### Competing Interest Statement

The authors have declared no competing interest.

