## Supplemental Figures for "Mammalian Milk Glycomes: Connecting the Dots between Evolutionary Conservation and Biosynthetic Pathways"

### Supplementary Figures

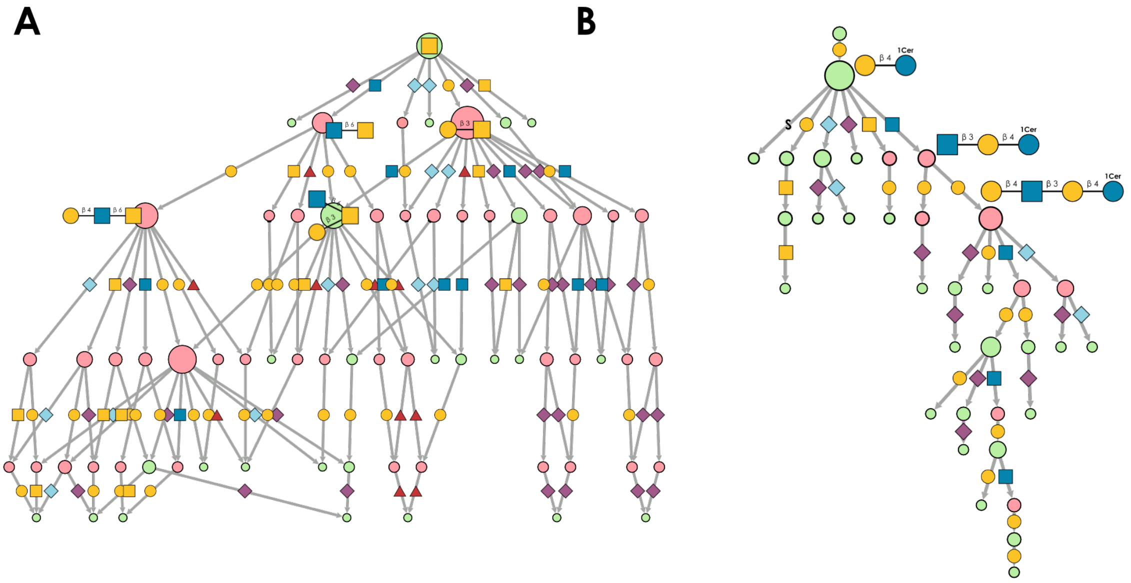


**Figure S1. Biosynthetic networks of other glycan classes; Related to Figure 1. (A-B)** We used all *O*-linked glycans from *Mus musculus* (mouse; A) and glycolipids from *Ovis aries* (sheep; B) that were contained in glycowork (version 0.6) to build unpruned biosynthetic networks with the construct_network function from glycowork. Inferred nodes are indicated in pink and observed nodes in green. The edge label indicates the type of added monosaccharide according to the SNFG. Node size is scaled by out-degree. Representative nodes are labeled. Networks were visualized in Cytoscape (version 3.9.1).

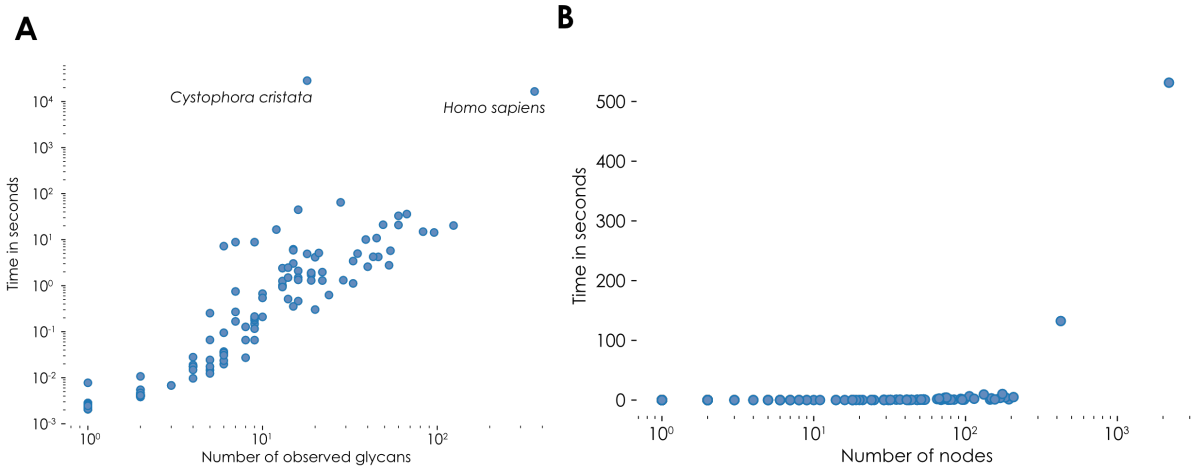


**Figure S2. Estimating speed of network construction and pruning; Related to Figure 1. (A-B)** For all species in our milk oligosaccharide dataset, we timed the construction (A) or pruning (B) of their biosynthetic network and plotted it against the number of glycans (A) or the size of the network (B). Computation was done using an Intel® Xeon® CPU @ 2.20GHz. The outlier in (A), *Cystophora cristata*, is caused by the presence of extremely long reported milk glycans coupled with the absence of reported medium-length structures, leading to a combinatorial explosion of possible pathways. In general, the length distribution for a species is much smoother and both network construction and pruning scale well with dataset size for realistic dataset sizes.

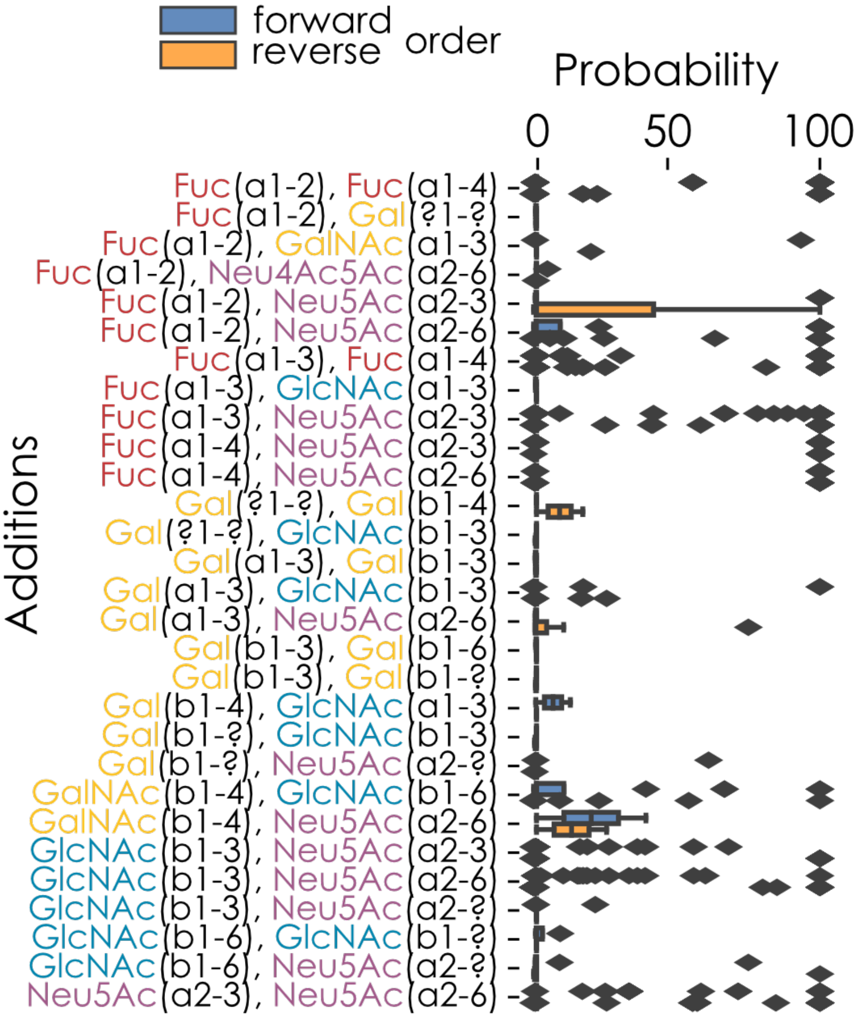


**Figure S3. Biosynthetic reactions showing no apparent path dependence; Related to Figure 2.** Similar to Figure 2E, for each observed addition of two monosaccharides in diamond-shaped network motifs, we grouped their probabilities across glycan sequence contexts (i.e., monosaccharides A and B could be added to glycan X, Y, Z, etc.) in both the forward (A-then-B) and reverse (B-then-A) order. Shown are the additions with a mean difference of below 15 with no significant path dependence, based on mean differences via two-tailed Welch's t-tests ($p > 0.05$). Data are depicted as mean values, with the box edges indicating the quartiles and the whiskers indicating the remaining data distribution.

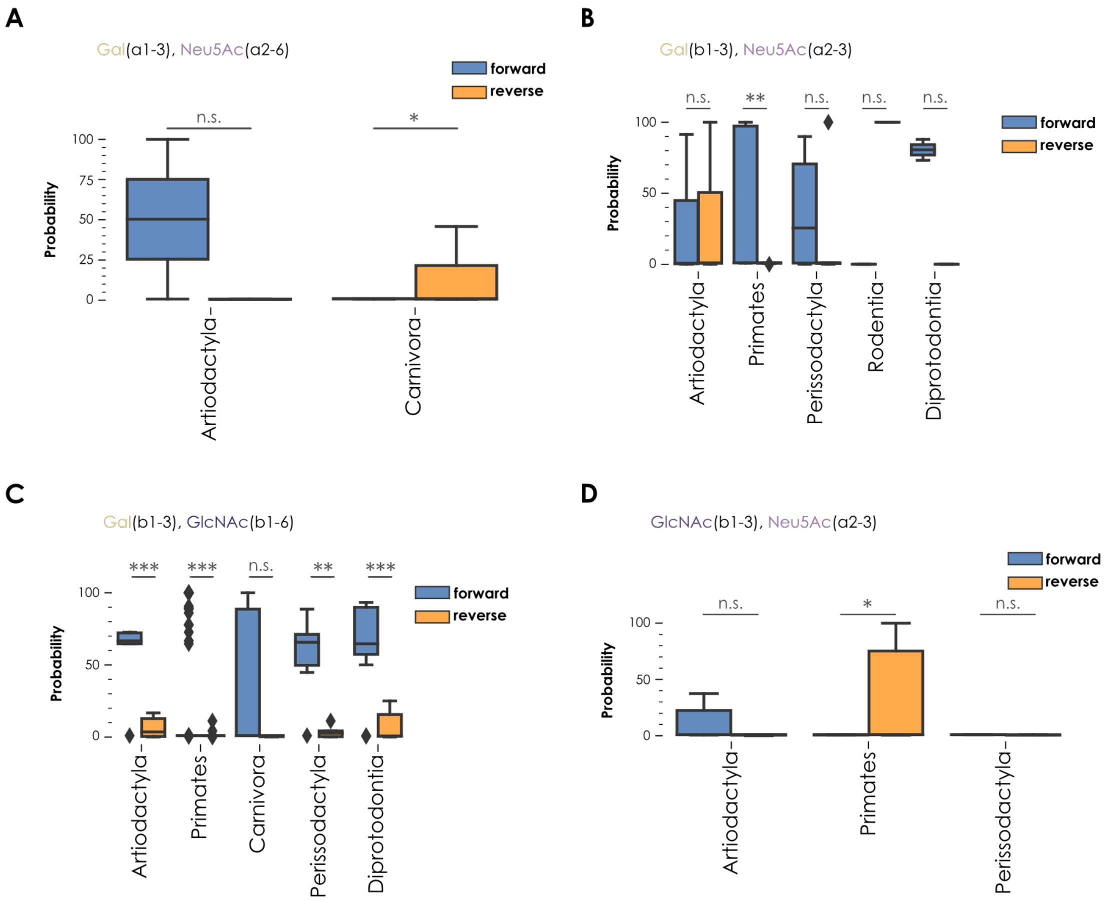


**Figure S4. Divergent path dependences across orders; Related to Figure 2. (A-D)** Similar to Figure 2E, for each observed addition of two monosaccharides in diamond-shaped network motifs, we computed and grouped their probabilities by taxonomic order (Artiodactyla, Perissodactyla, Diprotodontia, Primates, Carnivora, and Rodentia) across glycan sequence contexts in both the forward and reverse order. Shown are additions where at least one animal order displayed a strong divergence (i.e., the specific usage of the two possible path orders) compared to the others. Animal groups without any observation of a given diamond-shaped motif are not displayed. Artiodactyla and Carnivora showed an opposite order preference for adding Galα1-3 and Neu5Acα2-6 (A). Artiodactyla specifically performed the addition of Galβ1-3 and Neu5Acα2-3 (B) in both forward and reverse order. Similarly, Artiodactyla and Diprotodontia seemed to be able, although to different degrees, to conduct both forward and reverse addition of Galβ1-3 and GlcNAcβ1-6 (C). Similar to adding Galα1-3 and Neu5Acα2-6, addition of GlcNAcβ1-3 and Neu5Acα2-3 is performed in an opposite order between Artiodactyla and Primates (D). The few observations from Rodentia in (B) and Perissodactyla in (D) were not sufficient for any robust interpretations. Data are depicted as means, with box edges indicating quartiles and whiskers indicating the remaining data distribution. Mean difference was tested via two-tailed Welch's t-tests. ***, $p < 0.001$; **, $p < 0.01$; *, $p < 0.05$; n.s., $p > 0.05$.

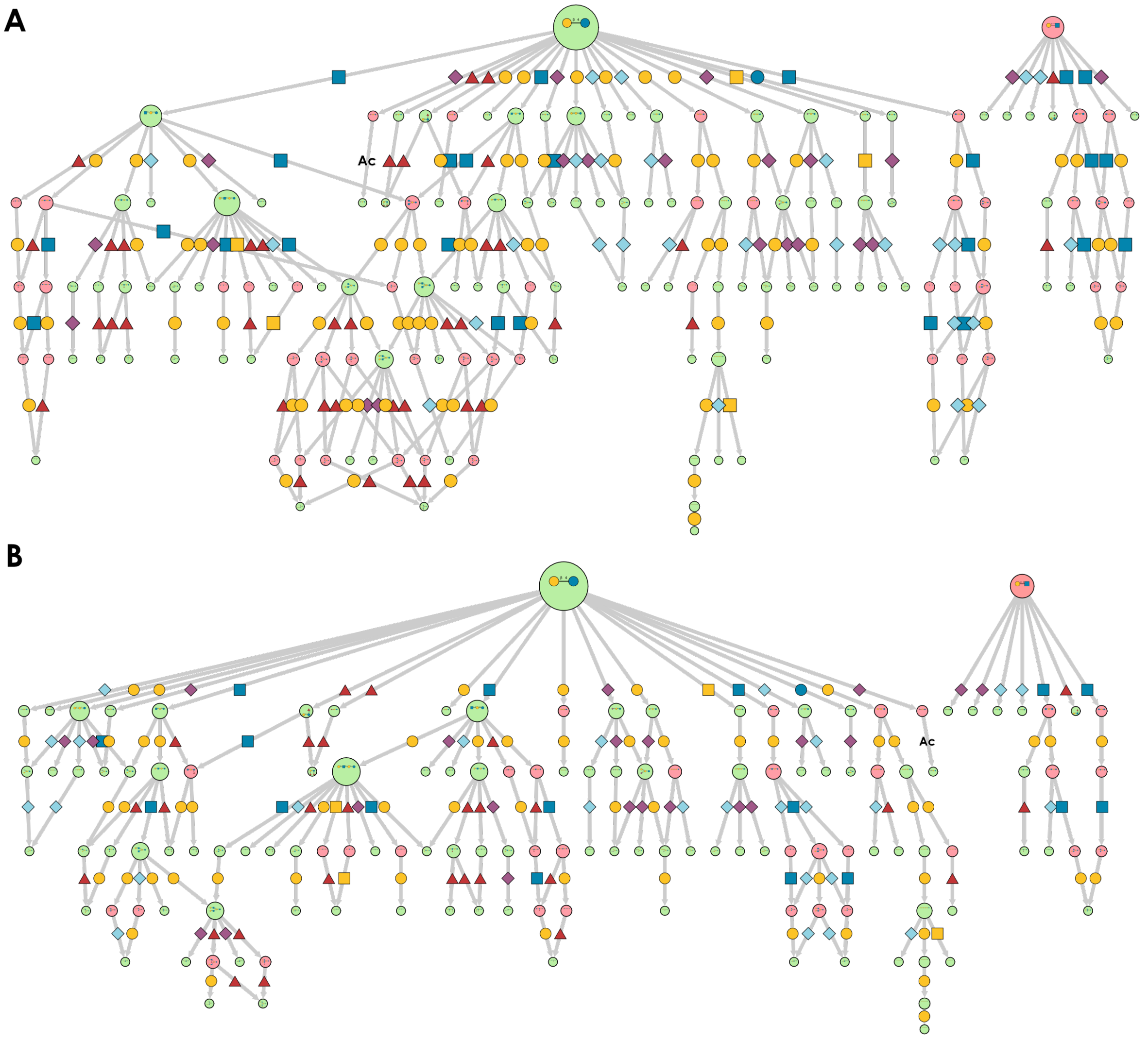


**Figure S5. Maximum-likelihood biosynthetic networks; Related to Figure 2. (A-B)** We constructed a biosynthetic network from goat (*Capra hircus*) milk oligosaccharides and pruned it (A). Then, we used the reaction path dependence described in Figure 2D to further specify the network (B). For this, we considered all still-remaining diamond-shape motifs and assessed whether any of their paths (i) had an absolute reaction path probability of above 50%, (ii) a probability more than 50% higher than the alternative path, and (iii) the removal of the alternative path did not abolish network connectivity. If all conditions applied, we removed the virtual node and its corresponding edges from the alternative path.

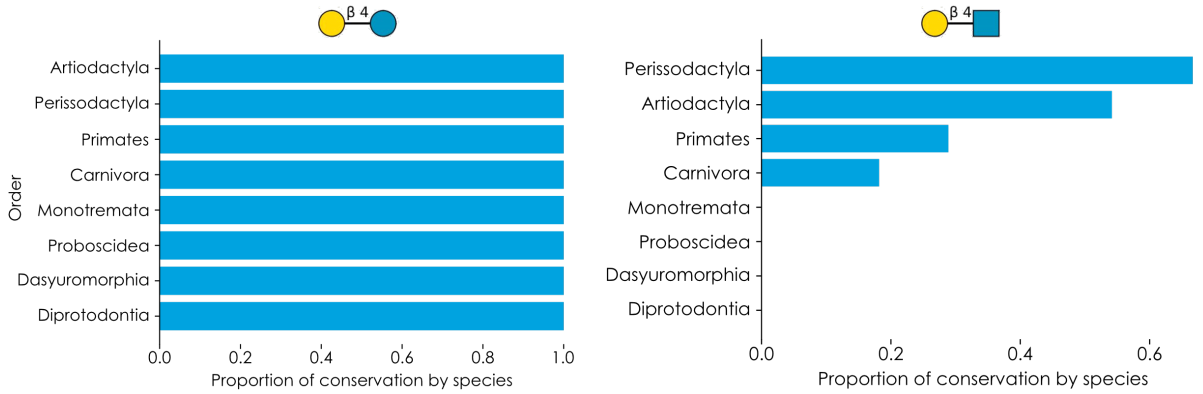


**Figure S6. Evolutionary conservation of lactose- and lactosamine-based milk glycans; Related to Figure 4.** For both lactose- and lactosamine-reducing ends, we used the check_conservation function in glycowork to estimate the evolutionary conservation across mammalian orders by assessing the proportion of species in each order that exhibited the indicated motif in their milk oligosaccharide biosynthetic networks.

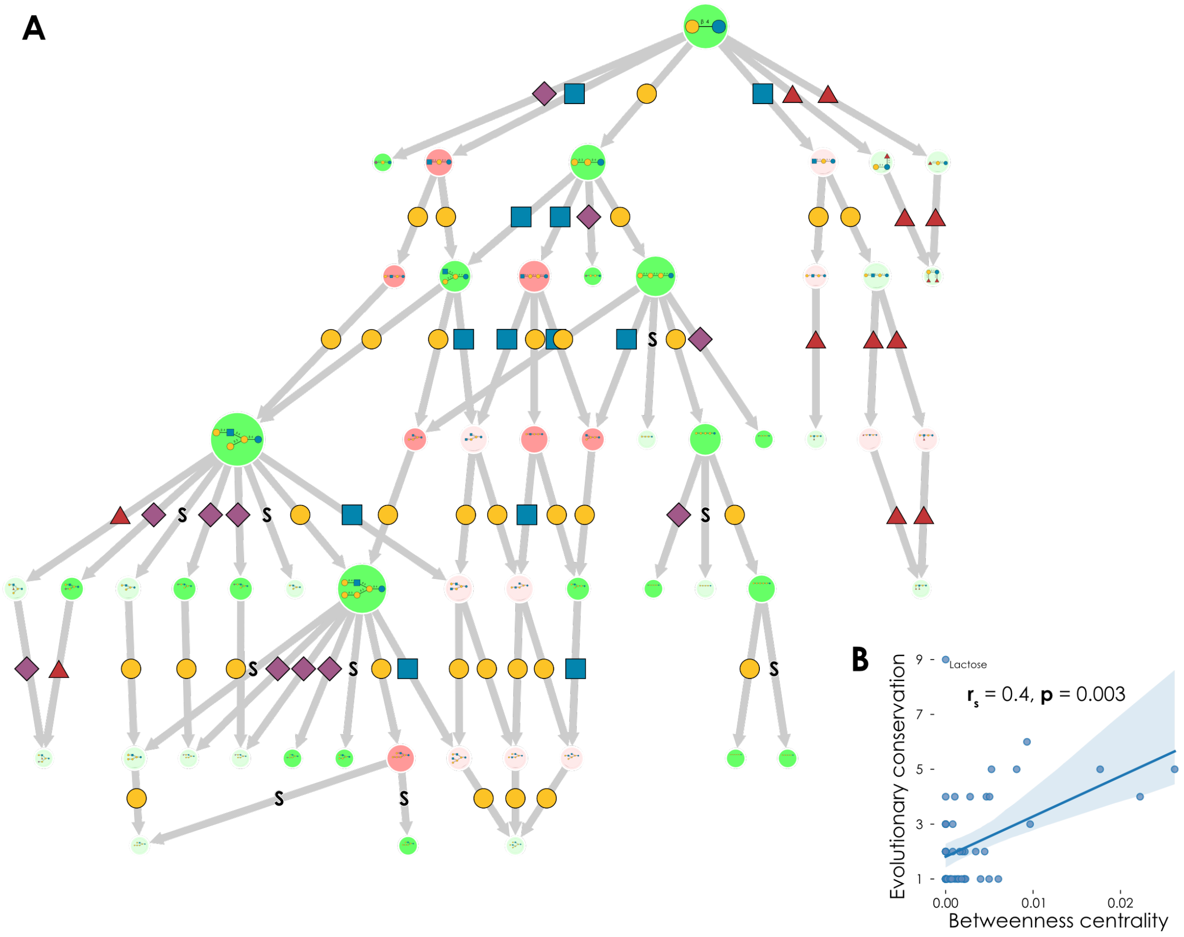


**Figure S7. Evolutionary network conservation across Diprotodontia; Related to Figure 4. (A)** Using the set of all milk oligosaccharides reported in the taxonomic order Diprotodontia, we constructed a biosynthetic network that we then evopruned. Using the highlight_network function from glycowork, we then estimated the evolutionary conservation of each glycan across Diprotodontia, visualized as increasing node transparency with decreasing conservation. Nodes that needed to be inferred in all species are indicated in pink and nodes that were observed in at least one species in green. The edge label indicates the type of added monosaccharide according to the SNFG. Node size is scaled by out-degree. Representative nodes are labeled. Networks were visualized in Cytoscape (version 3.9.1). **(B)** Calculating the Spearman rank correlation between glycan conservation (expressed as number of species in Diprotodontia exhibiting a given glycan, both inferred or observed) and betweenness centrality in the network in (A) showed that glycans are significantly more likely to be conserved across species if they are central to network connectivity.
